# Moisture Matters: Unintended Consequences of Performing Wet Sanitation in Dry Environments

**DOI:** 10.1101/2025.11.26.690674

**Authors:** Calvin Slaughter, Shihyu Chuang, Devin Daeschel, Lynne McLandsborough, Abigail B. Snyder

**Affiliations:** Department of Food Science, Cornell University, Ithaca, NY, USA; Department of Food Science, University of Massachusetts, Amherst, MA, USA

## Abstract

Cross-contamination of low-moisture foods (LMFs) with pathogens from equipment and environmental surfaces during production is a food safety concern. Wet sanitation is sometimes employed to mitigate cross-contamination in LMF facilities, but the introduction of moisture to otherwise dry environments can inadvertently promote pathogen growth. This study evaluated the risks associated with wet sanitation in LMF facilities by characterizing evaporation kinetics on powdered infant formula (PIF)–soiled surfaces, monitoring relative humidity (RH) in a LMF facility during and after wet sanitation, and assessing growth of *Salmonella*, *Listeria monocytogenes*, *Cronobacter sakazakii*, and *Enterococcus faecium* spot inoculated on PIF-soiled stainless steel coupons under dynamic RH conditions. As expected, higher RH slowed drying of PIF-soiled surfaces, prolonging periods when the soil’s water activity (*a*_w_) was high enough to support microbial growth. Correspondingly, all four organisms grew significantly at 81 and 97% RH over 120 h (p<0.05), while only *E. faecium* grew significantly below 81% RH (p<0.05). Monitoring of RH during and after wet sanitation in a commercial facility revealed spikes up to 100% RH during sanitation and sustained RH above 75% for more than 7 h in poorly ventilated areas. When those facility RH conditions were simulated in the laboratory, *Salmonella* populations on PIF-soiled coupons increased by more than 3.5 Log CFU/coupon within 66 h. These findings demonstrate the potential for wet sanitation to unintentionally enable environmental pathogen growth and highlight the importance of moisture and RH control in LMF facilities.

**Importance:** Wet sanitation is commonly employed by LMF manufacturers for allergen changeovers and to prevent cross-contamination from surfaces, but regulators, manufacturers, and researchers have all expressed concerns that wet sanitation may promote the growth of pathogens in otherwise dry production environments. Despite these concerns, research on the impact of wet sanitation on facility RH and its influence on microbial proliferation in low moisture production environments remains limited. This study provides evidence that wet sanitation substantially increases facility RH, leading to persistent hydration of soiled surfaces, creating conditions that enable microbial growth. These findings reinforce concerns over the use of wet sanitation in LMF production. This also demonstrates the need for reducing water use in LMF production facilities, implementing RH control strategies, as well as the adoption of alternative or supplemental dry sanitation strategies to mitigate microbial risks.

## Introduction

Low-moisture foods (LMFs) do not support the growth of infectious foodborne pathogens including *Salmonella*, *Listeria monocytogenes*, or *Cronobacter sakazakii*. *Salmonella* and *L. monocytogenes* are unable to grow below *a*_w_ 0.94 and 0.92, respectively (FDA, 2024). Based on the findings of Dancer et al. (2009), the minimum *a*_w_ for the growth of *C. sakazakii* is close to 0.94. Despite this innate barrier to infectious pathogen growth, between 2012 and 2020 LMFs were implicated in 54 confirmed outbreaks, and 1,659 product recalls due to pathogen contamination (Acuff et al., 2023). These events can be the result of product cross-contamination from equipment or environmental surfaces harboring pathogens (Beuchat et al., 2013). LMF processing environments are often dry, having low RH and minimal water usage. Under these dry conditions, pathogens that enter the facility through raw ingredients, personnel, or pests can persist on surfaces for weeks to months in dormant states (Bashir et al., 2022; Xie et al., 2024).

One of the few instances in which water is intentionally introduced into LMF facilities is during wet sanitation conducted to mitigate allergen cross-contact and the cross-contamination risks posed by these pathogens. Wet sanitation typically consists of the application of water and cleaning agents to remove soils and other objectionable matter from food contact and environmental surfaces, followed by the application of chemical sanitizers to chemically inactivate residual microorganisms (FDA, 2025). In practice, however, complete soil removal is relatively difficult to achieve across every single niche of the entire surface area of a large production facility. Factors such as poor equipment design (e.g., niches, crevices), surface roughness, and limited accessibility for cleaning tools hinder effective cleaning (Ihara et al., 2025). Moreover, surfaces that appear visually clean may still retain low levels of food residues, undetectable without biochemical assays (Daeschel et al., 2023; Fallahizadeh et al., 2025).

Food soils that are moistened but not removed by wet cleaning enhance food safety risk as they both shield pathogens from chemical sanitizers and are a nutrient-rich, high-moisture matrix conducive to microbial growth (Chowdhury et al., 2019; Cordier, 2014). Indeed, Cordier (2014) demonstrated that in the days following moisture introduction in a LMF facility, Enterobacteriaceae populations on surfaces increased dramatically. The potential of wet sanitation induced microbial growth was also highlighted in the U.S. Food and Drug Administration (FDA)’s draft guidance for LMF sanitation (FDA, 2025). In their draft guidance, the FDA recommends that water use during sanitation should be minimized and that surfaces should be dried promptly to prevent microbial growth. Recommended strategies to accelerate drying of surfaces in LMF facilities to prevent microbial growth include the hygienic design of equipment to promote drainage, application of forced-air systems, and the removal of standing water using squeegees or vacuums (Moerman & Mager, 2016).

Although the risks associated with wet sanitation and the importance of drying steps in low-moisture food (LMF) facilities are well recognized, quantitative data on how wet sanitation affects pathogen growth and persistence in these environments remains scarce yet necessary for estimating risk trade-offs between dry and wet sanitation strategies (Daeschel et al., 2025). Consequently, many food safety policies and practices are based on an incomplete understanding of how pathogens interact with moisture in LMF settings. This gap can limit the effectiveness of food safety strategies, where managing moisture to prevent pathogen growth may be just as critical as reducing initial contamination through sanitation. To address this need, we evaluated how moisture introduced during wet sanitation influences ambient RH, residual surface moisture, and subsequent pathogen growth.

## 3. Materials and Methods

### 3.1 Evaporation kinetics

A dynamic vapor sorption (DVS) analyzer (DVS Adventure, Surface Measurement Systems, United Kingdom) was used to assess the evaporation behavior of a rehydrated PIF matrix (2 g PIF mixed with 10 g of 0.1% phosphate buffered saline (PBS; Thermo Fisher Scientific, Waltham, MA, USA)) as it was exposed to controlled levels of RH. Samples (100 µg) were hung from a microbalance (0.1 µg resolution) on a stainless steel pan, inside of a humidity cell that was continuously purged with differential portions of dry (nitrogen) and wet (distilled, deionized water) gases to maintain a target vapor pressure, namely 33%, 51%, 64%, 81%, and 97% RH, at 22 ± 0.1°C, and at a volumetric flow rate of 20 standard cubic cm per min (cm^3^ ⋅ min^−1^), the instrumental lower limit to simulate little to no airflow. Positive and negative changes in mass reflected vapor adsorption and desorption, respectively. Data was acquired every other second. The PIF had a moisture content of 3.17 ± 0.02% w/w, as determined thermogravimetrically (MX-50, A&D Company, Ltd., Japan). Bulk Moisture Content of the PIF-PBS system was expressed in % wet basis, by subtracting the PIF’s solid content from the prevailing DVS mass reading, and dividing the difference by that same mass reading. Upon equilibrium, the moisture contents across the five RH conditions were plotted against the corresponding *a*_w_, which was assumed to be equivalent to the RH at which equilibrium was reached (RH/100). This relationship was used to generate a moisture sorption isotherm for the PIF matrix, characterizing its hygroscopic behavior from *a*_w_ 0.33 to 0.97. The resulting isotherm was then fit to a second-order polynomial model using R v4.5.0 (R Foundation for Statistical Computing, Vienna, Austria) in RStudio.2025.05 (Posit Software, PBC, Boston, MA).

### 3.2 Coupon Preparation

Coupons measuring 1.5 cm × 3.5 cm × 0.28 cm were cut from 2B-polished 304 stainless steel sheets originally sized 30 cm × 30 cm × 0.28 cm. Coupons were sterilized at 121°C for 30 min. Control coupons were set aside immediately following sterilization. Remaining coupons were then soiled with 0.02 ± 0.003 g PIF (Similac Total Comfort, Abbott Nutrition, Columbus, OH, USA) spread evenly across the surface of the coupon using a sterile 10 μL inoculation loop. The mass of the PIF was measured using an Aczet CY 314 C analytical balance (Maharashtra, Mumbai, India). PIF was selected as the soil of interest for this study due to its high nutrient density and low background microbial load.

### 3.3 Culture Growth and Inoculation

Three-strain cocktails of *Salmonella enterica* (FSL M8-0476, FSL M8-0481, FSL R9-5220), *Listeria monocytogenes* (FSL F2-0941, FSL F3-0842, FSL L4-0253), and *Cronobacter sakazakii* (FSL F6-0024, FSL F6-0032, FSL F6-0046) were used in this study. The *Salmonella* and *C. sakazakii* strains used were isolated from low moisture foods (LMFs). The *L. monocytogenes* isolates used were environmental isolates from food processing facilities. In addition to the above pathogens, *Enterococcus faecium* NRRL B-2354 was included due to its common use as a surrogate for *Salmonella* in LMF safety research (Bianchini et al., 2014; L. Chen & Snyder, 2023; S. Liu et al., 2018).

Strains were cultured using the methods described by Hildebrandt et al. (2016), with minor modifications. Frozen cultures stored in Tryptic Soy Broth (TSB; Thermo Fisher Scientific, Waltham, MA, USA) at −80 °C in 20% glycerol were inoculated into 10 mL of brain heart infusion (BHI; Thermo Fisher Scientific, Waltham, MA, USA) broth using sterile 1 µL inoculation loops. Cultures were incubated at 37°C for 24 h, followed by a 1 µL transfer into fresh 10 mL BHI broth using a sterile 1 uL inoculation loop and a second 24 h incubation at 37°C. Following the second incubation, 300 µL of each culture was spread plated across three 100 mm × 15 mm plates of BHI agar and then incubated at 37°C for 24 h. Strains were grown on agar instead of liquid media to promote desiccation resistance associated with dry growth conditions typical of LMF environments (Hildebrandt et al., 2016).

After incubation, bacterial cells from each strain’s three agar plates were harvested in 10 mL of 0.1% PBS using sterile 10 µL inoculation loops. Cultures were centrifuged at 3,000 × *g* for 15 min (Eppendorf 5804R, Eppendorf, NY, USA), the supernatant discarded, and the pellet resuspended in 10 mL PBS to yield approximately 10⁹ CFU/mL. Concentrated three-strain cocktails were prepared by combining 1 mL of each strain suspension in PBS and homogenizing the mixture using a vortex mixer. Concentrated cocktails were then serially diluted in PBS to a working concentration of approximately 10⁴ CFU/mL.

For inoculation, 100 µL of each diluted 3-strain cocktail was pipetted onto the center of a PIF-soiled stainless steel coupon, yielding approximately 3 Log (CFU)/coupon. The inoculum was evenly spread across the surface using sterile 10 µL inoculation loops. Negative growth controls were prepared by pipetting 100 µL of 3-strain cocktail onto non-soiled coupons and spreading the liquid evenly using sterile 10 µL inoculation loops.

### 3.4 Static RH Growth Curves for Salmonella, Listeria monocytogenes, C. sakazakii, and E. faecium

RHs of 33%, 51%, 64%, 81%, and 97% were established in glass desiccator chambers maintained at approximately 22°C using saturated salt solutions of magnesium chloride, magnesium nitrate, sodium nitrite, ammonium sulfate, and potassium sulfate, respectively. Initial RH levels in each desiccator were measured after a 48 h equilibration period using a RH meter (OM-EL-WIFI-T, DwyerOmega, Michigan City, IN, USA), with subsequent verifications performed at 24 h intervals.

Soiled coupons inoculated with *E. faecium* or 3-strain cocktail of *Salmonella*, *L. monocytogenes*, or *C. sakazakii* were placed on plastic racks within the desiccators. At predetermined time points (0, 24, 48, 72, 96, and 120 h), coupons were removed for microbial enumeration. Growth curves for each organism were constructed by plotting the average Log (CFU)/coupon across four biological replicates against time.

### 3.5 Facility RH Data Collection

Changes in RH during and after routine wet sanitation were investigated within a 48 h period in two dry-blending lines within a large-scale spice mixing facility. Each dry blending line consisted of two levels separated by concrete flooring with sections cut out for equipment to pass through: the upper level housed a ribbon blender, while the lower level contained a hopper–auger filler and conveyor system. Line 1 was equipped with a 3,000 lb ribbon blender, whereas Line 2 housed a smaller 500 lb ribbon blender. The dimensions of the upper and lower levels were approximately 20 ft × 30 ft × 8 ft and 20 ft × 30 ft × 10 ft, respectively (Figure 3).

Wet sanitation of the dry blending lines was conducted approximately once daily, either between production lots or during product changeovers. The sanitation process included an initial rinse of equipment and environmental surfaces using water hoses, followed by application of a detergent cleaner. Following addition of the detergent, the mixers were filled with water and allowed to run for 0.9-1.6 h. This was followed by a secondary water rinse of both equipment and environmental surfaces, and ATP swabbing of equipment surfaces to verify cleaning effectiveness. Finally, an aqueous quaternary ammonium compound-based sanitizer was applied to equipment and environmental surfaces using spray hoses. Post-sanitation drying conditions varied between the two lines. For Line 1, the doors to both the upper and lower rooms were opened, and compressed air was used to manually dry environmental surfaces prior to the start of production. In contrast, Line 2 remained sealed, with doors closed and no use of compressed air for drying.

During the sanitation process, RH was recorded at 1 s intervals using handheld RH loggers (HOBO MX2302 External Temp/RH data logger, LI-COR, Lincoln, NE, USA) positioned on the upper level of each dry blending line. Handheld devices were used to prevent water damage during cleaning. Once sanitation was completed, RH loggers were fixed in place at the return vent, upper door, lower double-door, lower door and the roller conveyor (Figure 3). RH data was collected in 1 s intervals through the end of the subsequent production cycle.

### 3.6 Growth Curves Under Simulated Facility Conditions

RHs of 33%, 51%, 64%, 76%, and 97% were established in glass desiccator chambers maintained at approximately 22°C using saturated salt solutions of magnesium chloride, magnesium nitrate, sodium nitrite, sodium chloride, and potassium sulfate, respectively. Initial RH levels in each desiccator were measured after a 48 h equilibration period using RH meters (OM-EL-WIFI-T, DwyerOmega, Michigan City, IN, USA), with subsequent verifications performed at 24 h intervals.

Coupons prepared and inoculated as described in Sections 3.2 and 3.3 were subjected to two RH exposure conditions: a dynamic treatment simulating RH recorded during and after wet sanitation in a commercial facility, and static low-RH treatment. The dynamic RH treatment consisted of three sequential humidity phases: 30 min in the 97% RH desiccator, 9 h in the 76% RH desiccator, 9 h in the 64% RH desiccator, and 5.5 h in the 51% RH desiccator, cycled continuously for a total duration of 72 h. These conditions were selected based on RH measurements recorded in a commercial facility during sanitation and post-sanitation at the upper door location, where persistently high RH levels were observed. The static treatment consisted of constant exposure to 33% RH, representing conditions under which humidity is tightly controlled.

For both conditions, coupons were re-wetted at 24 and 48 h with 100 µL of sterile PBS, evenly spread across the surface using sterile 10 µL inoculation loops. Re-wetting was incorporated to simulate daily wet sanitation practices observed in the facility. Coupons were removed for enumeration at 0, 9.5, 18.5, 24, 33.5, 42.5, 48, 57.5, 66.5, and 72 h, as described in Section 3.7.

### 3.7 Microbial Enumeration

Coupons were removed from desiccators and placed into sterile 3″ × 5″ (7.6 × 12.7 cm) Whirl-Pak® bags (Whirl-Pak, Madison, WI, USA) containing 10 mL of 0.1% buffered peptone water (BPW). Each coupon was manually stomached for 30 s to dislodge PIF and bacteria and homogenize them into the BPW. From the resulting BPW suspension, serial dilutions ranging from 10⁻² to 10⁻⁶ were prepared.

For enumeration, 500 µL of the undiluted BPW suspension was plated across five 100 mm × 15 mm BHI agar plates (100 µL per plate), and 100 µL of each serial dilution was spread-plated onto individual BHI agar plates. Plates were incubated at 37°C for 24 h. Following incubation, colony-forming units (CFU) per coupon were calculated based on colony counts on the BHI plates.

### 3.8 Statistical Analysis

Statistical comparisons of bacterial populations at sampling intervals as compared to the time of inoculation were done using two sample student-t tests with a subsequent Benjamini-Hochberg adjustment to control for familywise error rate (Chen et al., 2017). Results were deemed significant if their p-value was below 0.05. All statistical analyses were conducted using R version 4.5.1 (R Foundation for Statistical Computing, Vienna, Austria) in RStudio.2025.05 (Posit Software, PBC, Boston, MA).

## 4. Results

### 4.1 High RH Slowed the Drying of Hydrated Food Soil

The drying kinetics of hydrated PIF soil were examined across RHs ranging from 33% to 97%. Drying proceeded through distinct constant- and falling-rate stages driven by moisture gradients within the soil and between the soil surface and the surrounding environment.

Both the initial drying rate and the duration of the constant and falling rate periods were strongly influenced by RH. As expected, lower RH conditions accelerated drying, resulting in higher initial drying rates and shorter drying phases. The highest initial drying rate (~0.13 mg·min⁻¹) was observed at 33% RH. The constant rate period at 33% RH lasted approximately 8 h, followed by falling rate period I (8–13 h) and falling rate period II (13–20 h). As RH increased, drying slowed progressively. At 51% RH, the constant rate (~0.10 mg·min⁻¹) persisted for 12 h, followed by falling rate period I (12–15 h) and falling rate period II (15–24 h). At 64% RH, the constant rate (~0.09 mg·min⁻¹) lasted 15 h, with falling rate period I from 15 to 17.5 h and falling rate period II from 17.5 to 26 h. At 81% RH, the constant rate (~0.07 mg·min⁻¹) extended to 20 h, followed by falling rate period I (20–23.5 h) and falling rate period II (23.5–28 h). The slowest drying occurred at 97% RH, where the constant rate (~0.05 mg·min⁻¹) persisted for approximately 25 h before transitioning into falling rate period I (25–32 h) and falling rate period II (32–39 h) (Figure 1A).

**Figure 1.**
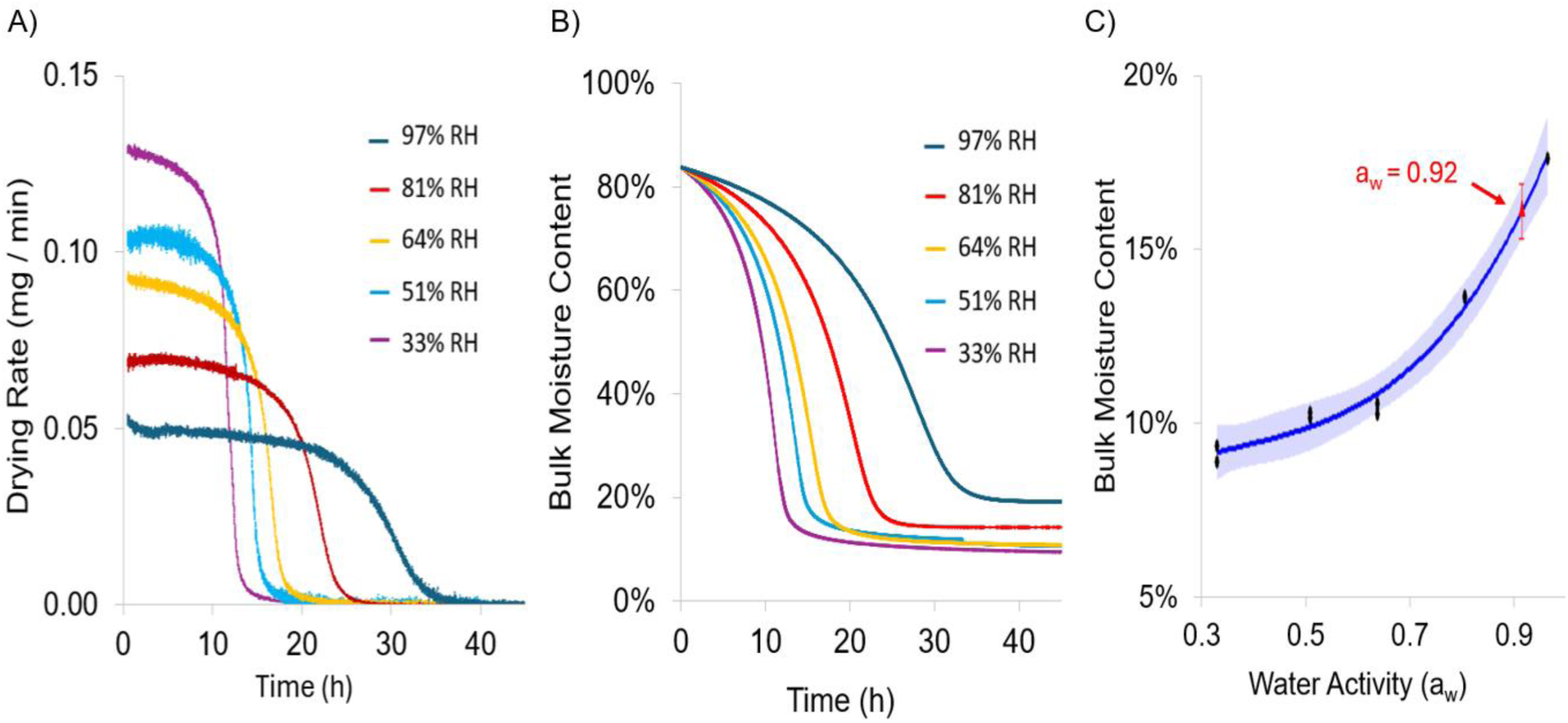
Average drying rate (mg/min) of PIF soil at 97, 81, 64, 51, 33% RH over 2 replicates **(A)**. Average bulk moisture content over time of PIF soil during drying at 97, 81, 64, 51, 33% RH over 2 replicates **(B)**. Moisture de-sorption isotherm of PIF soil with a 95% confidence interval **(C).**

The observed drying trends reflect the increasing resistance to moisture removal at higher RH, where a reduced vapor pressure gradient between the soil and the environment slows both surface evaporation and internal diffusion. Moisture loss from soils during constant-rate periods is dominated by the removal of free surface water through evaporation (Richardson et al., 2002). During falling-rate period I, surface evaporation becomes limited by the diffusion of liquid water from the soil interior to the exterior, causing a progressive decline in the drying rate (Richardson et al., 2002). Similarly, during the second falling-rate period, drying at the surface is limited, resulting in a slow, vapor-phase diffusion of residual moisture from within the soil matrix (Richardson et al., 2002).

At the end of falling rate Period II, average equilibrium moisture content (EMC) across 2 replicates increased with RH, averaging 9.4, 10.6, 13.8, 14.2, and 19.1% EMC at 33% RH (*a*_w_ 0.33), 51% RH (*a*_w_ 0.51), 64% RH (*a*_w_ 0.64), 81% RH (*a*_w_ 0.81), and 97% RH (*a*_w_ 0.97) respectively (Figure 1B; Figure 1C).

To further characterize the relationship between *a*_w_ and bulk moisture content, a partial moisture desorption isotherm was constructed using the EMC values, enabling prediction of *a*_w_ from bulk moisture content across a broader range of RH (Figure 1C). Moisture content exhibited a positive, nonlinear relationship with *a*_w_, which was modeled using a second-degree polynomial regression (R² = 0.98). Although the moisture sorption isotherm constructed did not cover *a*_w_ below 0.33, or above 0.97, the region between these extremes was consistent with previously constructed moisture isotherms for PIF (Tham et al., 2016). From the constructed model, a *a*_w_ of 0.92, the lowest *a*_w_ supporting the growth of pathogens used in this study, corresponded to a moisture content of 16.89 ± 0.82%. The time required for the PIF soil to dry below this bulk moisture content varied with RH. It took 12.6 h to drop below *a*_w_ 0.92 at 33% RH, 15.6 h at 51% RH, 17.4 h at 64% RH, and 23.7 h at 81% RH (Figure 1B). For 33, 51, 64, and 81% RH, the transition below 16.89% moisture content occurred during falling rate periods. However, since falling rate periods are characterized by moisture gradients between the soil surface and interior, internal regions of the soil likely retained local moisture contents above 16.89% and *a*_w_ above 0.92 for a short time beyond these timepoints. As expected, at 97% RH, the soil never reached a moisture content of 16.89%, indicating it maintained a growth supporting *a*_w_ through the drying process, and at equilibrium (Figure 1B).

### 4.2 Significant Growth of Pathogens Occurred in Hydrated PIF at Elevated RH

Increased RH strongly enhanced microbial survival and growth on PIF-soiled coupons. Across all RH levels, no growth was observed on any of the non-soiled coupons. In contrast, PIF-soiled coupons at 97% RH supported growth and persistence of all four organisms (Figure 2). *E. faecium* and *C. sakazakii* populations increased significantly (p<0.05) by 24 h (+4.5 ± 0.3 and +2.7 ± 0.7 Log (CFU)/coupon, respectively), while *Salmonella* and *L. monocytogenes* grew significantly (p<0.05) by 48 h (+4.2 ± 0.8 and +3.8 ± 1.5 Log (CFU)/coupon, respectively). Populations of all four organisms remained elevated above 7 Log (CFU)/coupon through 120 h. These outcomes align with the drying data indicating that at 97% RH the soil never reached *a*_w_ 0.92 (Figure 1B; Figure 2), thus consistently maintaining conditions favorable for growth and persistence.

**Figure 2.**
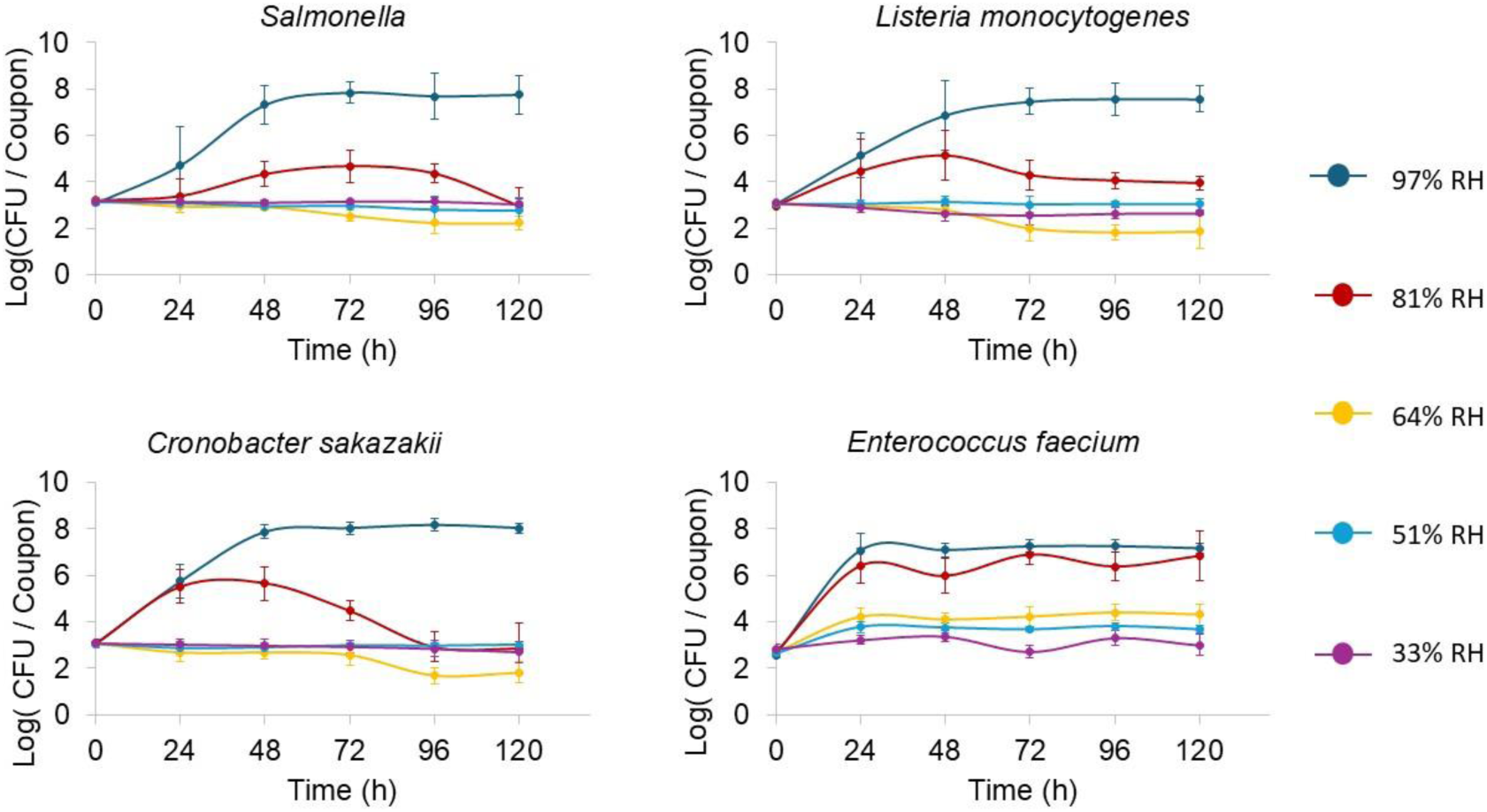
Growth curves of *Salmonella* **(Top-left)**, *Listeria monocytogenes* **(Top-right)**, *Cronobacter sakazakii* **(Bottom-left)**, and *Enterococcus faecium* **(Bottom-right)**, on PIF-soiled stainless steel coupons incubated at 22°C at 97%, 81%, 64%, 51%, and 33% RH over 120 h. Error bars represent the standard deviation of biological replicates (n= 4).

At 81% RH, all four organisms exhibited significant growth on soiled coupons, though the extent varied significantly by organism (Figure 2). *E. faecium* and *C. sakazakii* showed significant increases in population by 24 h (p<0.05), with population gains of +3.7 ± 0.5 and +2.4 ± 0.7 Log (CFU)/coupon, respectively. Although the PIF soil was predicted to drop below *a*_w_ 0.92 by 23.4 h, *Salmonella* and *L. monocytogenes* did not exhibit statistically significant growth (p > 0.05) until after the 24 h mark. *Salmonella* increased significantly by 48 h (p < 0.05) (+1.1 ± 0.5 Log (CFU)/coupon). *L. monocytogenes* displayed highly variable growth at earlier time points, with statistically significant growth observed only at 96 h (p < 0.05) (+1.1 ± 0.4 Log (CFU)/coupon). The continued growth of *Salmonella* and *L. monocytogenes* after the soil was expected to fall below the growth-enabling *a*_w_ of 0.92 is likely due to uneven moisture distribution during falling-rate drying periods. Notably, the population of *Salmonella*, *L. monocytogenes*, and *C. sakazakii* gradually decreased over time following their initial growth, a trend not observed at 97% RH. This difference reflects the difference in PIF soil moisture between these two conditions. At 97% RH, the soil retains enough moisture to support microbial growth, allowing bacterial populations to replenish as cells die. In contrast, at 81% RH, the soil’s moisture content became too low to sustain growth, so dead bacterial cells are not replaced, leading to a gradual decline in population (Bashir et al., 2022).

At 64% RH, *Salmonella*, *C. sakazakii*, and *L. monocytogenes* populations did not increase on PIF-soiled coupons (Figure 2). Instead, populations declined gradually, with significant reductions of *C. sakazakii* at 96 and 120 h (−1.2 ± 0.4 and −1.2 ± 0.8 Log (CFU)/coupon, respectively) and for *L. monocytogenes* at 96 h (−1.2 ± 0.4 Log (CFU)/coupon). In contrast, *E. faecium* grew significantly, increasing by +1.5 ± 0.3 Log (CFU)/coupon by 24 h and remaining above 4 Log (CFU)/coupon through 120 h. Based on the drying kinetics of the PIF soil, the soil would have reached a *a*_w_ of 0.92 by ~17.3 h. For *Salmonella*, *L. monocytogenes*, and *C. sakazakii*, this relatively quicker drop below *a*_w_ 0.92 was sufficient to prevent significant growth. *E. faecium*’s growth at 64% RH suggests it can either grow at lower *a*_w_ than the other organisms, increase in population quickly enough to achieve significant growth before *a*_w_ drops below its growth threshold, or both. Additional evidence for the faster growth of *E. faecium* comes from its growth behaviors at high RH relative to the other organisms. At 97% RH, *E. faecium* increased by 4.5 ± 0.3 Log (CFU)/coupon within 24 h, whereas the other species grew less than 3 Log (CFU)/coupon in the same period.

At 51% and 33% RH, drying occurred rapidly (reaching 16.89% moisture content in 16.3 h and 12.4 h, respectively), leaving minimal time for microbial growth on the soiled coupons (Figure 2). Under these conditions, populations of *Salmonella*, *L. monocytogenes*, and *C. sakazakii* remained stable through 120 h, showing no statistically significant increases or decreases (p > 0.05). As observed at 64% RH, only *E. faecium* exhibited statistically significant growth, with modest increases at 51% RH (+1.0 ± 0.4 and +1.0 ± 0.3 at 72 and 120 h) and a small gain at 33% RH by 96 h (+0.5 ± 0.1). While *E. faecium* is a reliable surrogate for *Salmonella* in thermal inactivation studies (Bianchini et al., 2014; S. Liu et al., 2018), its consistently divergent growth behavior under varying RH conditions suggests it is an unsuitable surrogate for *Salmonella* for studies where microbial growth is a key consideration.

### 4.3 Wet Sanitation Resulted in Persistent Increases in RH Both During and After Sanitation

RH measurements recorded during and after sanitation of two high-hygiene areas within a LMF facility demonstrated that wet sanitation drastically increases RH, both during and after sanitation, and that intra-line RH measurements were heterogeneous by location.

During cleaning of Line 1’s blender (0.0–0.9h), RH varied between 50 and 65%. RH then peaked at 100% during the post-cleaning water rinse (0.9–1.5 h) before decreasing to between 60 and 75% as the quality team conducted post-cleaning adenosine triphosphate (ATP) and allergen swabs to verify cleaning efficacy prior to sanitizer application (1.5–3.1 h). Sanitizer application (3.1–3.6 h) also resulted in a noticeable spike in RH up to 88% (Figure 3).

**Figure 3.**
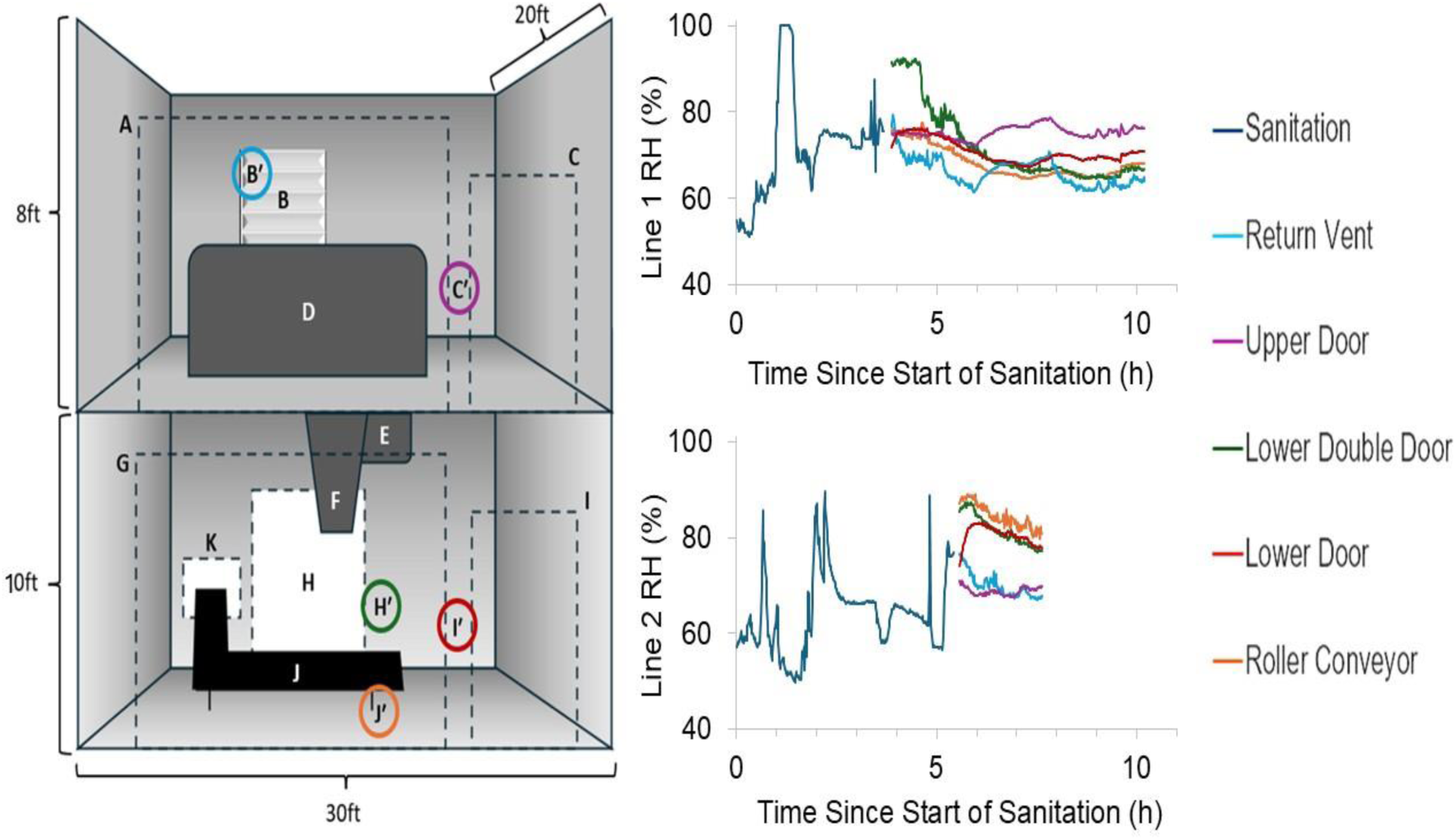
Schematic of the dry blending lines in which facility RH values were collected. (A) Upper Roller Door, (B) Return Vent, (B′) RH Logger, (C) Upper Door, (C′) RH Logger, (D) Dry Blend Mixer, (E) Auger Conveyor, (F) Hopper, (G) Lower Roller Door, (H) Lower Double Door, (H′) RH Logger, (I) Lower Door, (I′) RH Logger, (J) Roller Conveyor, (J′) RH Logger, (K) Roller Conveyor Exit **(Left)**. Recorded RH during sanitation of dry blending Lines 1 and 2, and after sanitation at multiple locations throughout the Lines **(Right)**.

Following sanitation, doors to both the upper and lower rooms of Line 1 were opened, and compressed air was used to manually dry environmental surfaces prior to the start of production. RH varied by location with no clear stratification between the upper and lower rooms. Over 6.3 h, RH at the air return vent, lower door, and roller conveyor locations ranged between 62 and 78%, with a slight downward trend over time. The lower double door initially started at 91% RH and persisted at this level for 0.7 h before gradually decreasing over 1.4 h to 64–68% RH through the end of data collection. The upper door location experienced minimal change in RH which fluctuated between 75 and 79% over the entire 7.1 h monitoring period, (Figure 3).

During the sanitation of Line 2, RH spiked to 86% during cleaning (0.0–1.6 h), and further increased during the post-cleaning water rinse (1.6–2.5 h), reaching 87% at 2.0 h and 90% at 2.2 h. Humidity then fell to between 68–58% as the quality team conducted post-cleaning swabs (2.5 h–4.7 h). During sanitizer application (4.7–5.4 h), RH spiked again to 88% at 4.8 h and 78% at 5.3 h. Unlike Line 1, the doors to both the upper and lower levels of Line 2 remained closed, and no compressed air was used to aid surface drying. This resulted in clear vertical stratification of RH, with consistently higher levels recorded in the lower room (Figure 3).

Immediately following sanitation, RH at the roller conveyor and lower double door locations of Line 2 fluctuated between 84–88%, gradually decreasing to 77–80% over 2.3 h. At the lower door, RH rose from 76% to 83% before declining back to 76% over 1.7 h. The return vent showed a steady decrease from 75% to 67% RH. Similar to Line 1, the upper door location in Line 2 exhibited minimal RH change, fluctuating between 68–71% throughout the 2.3 h post-sanitation period (Figure 3).

Differences in RH patterns between the two dry blending lines likely reflect variation in equipment size and post-sanitation drying practices. Line 1 contained a larger blender, which likely required more water during cleaning and sanitizing, leading to higher RH levels during sanitation. The combination of open doors and compressed air in Line 1 would have enhanced convective drying and air exchange, promoting uniform RH decline and preventing the vertical stratification observed in Line 2. In contrast, limited air movement on Line 2 could have contributed to the observed stratification between the upper and lower levels.

While RH stratification on Line 2 due to a lack of air circulation was to be expected, it was somewhat unexpected that the lower level of Line 2 had higher RH than the upper level. Since humid air at a given temperature is less dense and typically rises, we would expect the more humid air to accumulate at the upper level rather than the lower. Consistent with this principle, Song et al. (2019), reported that upper floors of residential buildings often exhibit elevated humidity levels relative to lower floors. A potential explanation for this discrepancy is that water from the upper level drained into the lower level, a factor not present in residential buildings. This drainage would result in increased surface moisture and evaporation at the lower level, thereby elevating the local RH.

The stable RH plateaus observed at the upper door location of Lines 1 and 2 suggest constraints that hinder moisture removal, creating potential niches for microbial persistence. Poor air circulation at the upper door location is a likely culprit for the observed plateau in RH. Indeed, Liu et al. (2024), found that areas within constructed environments with limited airflow retain moisture, resulting in elevated RH over long periods of time. Increasing ventilation in these areas was found to significantly reduce RH (Liu et al., 2024). However, even when deliberate steps to promote air flow were taken for Line 1, including the opening of doors and the use of air hoses, this elevated RH plateau persisted. This indicates these interventions alone may be insufficient to control post-sanitation humidity. Additional strategies such as portable dehumidifiers, improved ventilation design, or HVAC adjustments may be needed to achieve more rapid drying to reduce microbial risk.

### 4.4 *Salmonella* on PIF-soiled Coupons Grew Significantly Under Simulated Facility RH Conditions

Under simulated facility conditions, *Salmonella* populations increased by 0.5 ± 0.2 Log(CFU)/coupon within 18 h, a statistically significant rise from baseline (p < 0.01). Growth continued, peaking at 7.5 Log(CFU)/coupon by 66 h, followed by a slight decline to 6.9 Log(CFU)/coupon at 72 h. At 33% RH with re-wetting, *Salmonella* populations grew more slowly and reached lower levels. Statistically significant growth (p < 0.05) was not observed until 42 h (+0.5 ± 0.2 Log(CFU)/coupon). From 42 to 72 h, populations continued to increase, reaching an average of 4.5 Log(CFU)/coupon at 72 h.

Comparisons of trials at 33% RH with and without re-wetting (Figure 2) indicate that while strict RH control can suppress growth following isolated wetting events, it is insufficient when surfaces are repeatedly exposed to moisture, as occurs during frequent wet sanitation (Figure 2; Figure 4). This is consistent with Furtado et al. (2023) which found that desiccated *Salmonella* can rapidly resume growth once moisture is reintroduced, particularly after brief desiccation periods.

**Figure 4.**
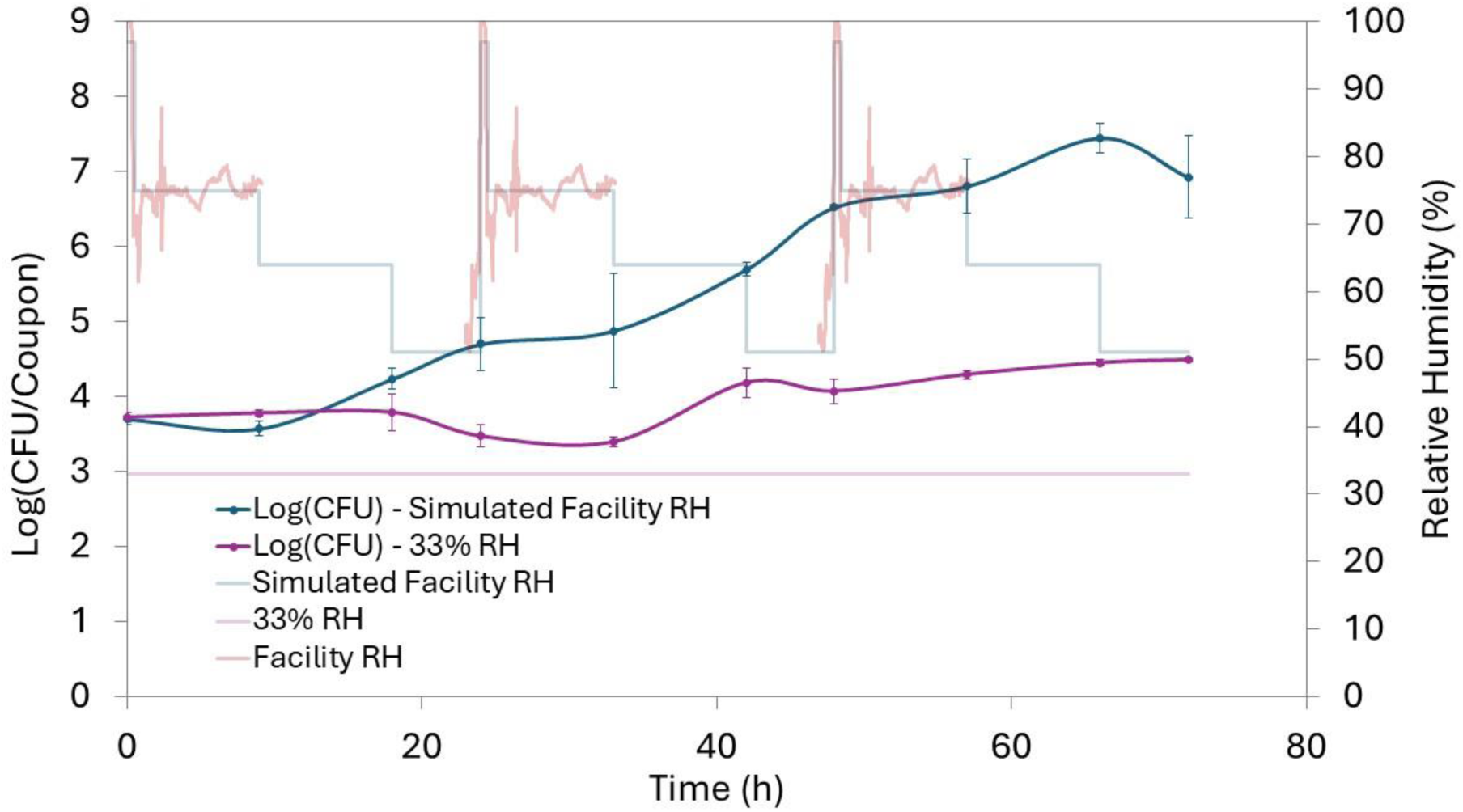
Population of *Salmonella* on PIF–soiled stainless steel coupons incubated at 22°C over 72 h under two RH conditions: constant RH at 33% (purple) and a dynamic RH profile (dark blue). The dynamic RH profile (light blue) simulates RH fluctuations recorded during and after sanitation of a dry blending line in a LMF facility (red). The constant 33% RH (light purple) represents a hypothetical scenario in which ambient RH is tightly controlled. Error bars represent the standard deviation of biological replicates (n = 3).

The difference in the magnitude of *Salmonella* growth between the simulated facility conditions and the static 33% RH demonstrates the role even temporarily elevated RH plays in enabling microbial growth on soiled surfaces. While the importance of minimizing water used during sanitation is well recognized, the importance of RH control in LMF facilities remains underemphasized in current LMF safety frameworks. In its *Code of Hygienic Practice for Powdered Formulae for Infants and Young Children*, the World Health Organization (WHO) acknowledges the risk of microbial growth following wet sanitation and emphasizes the importance of rapid drying after moisture introduction. The guidance also outlines tools and practices to facilitate and maintain a dry processing environment. The WHO’s recommended moisture control strategies include clean-out-of-place (COP) rooms, dry cleaning tools, and installing hygienically designed dry drains (*CXC 66*, 2008). While all these strategies contribute to keeping production environments dry and thus preventing microbial growth, the control and monitoring of RH are notably absent from the WHO’s guidance. Similarly, in its *2025 Draft Guidance for Establishing Sanitation Programs for Low-Moisture Ready-to-Eat Human Foods*, the FDA recommends that water introduced during sanitation be removed “as soon as possible” and that dry cleaning tools be used in place of wet sanitation whenever feasible. However, the guidance again omits any discussion of controlling RH following wet sanitation to prevent microbial growth (FDA, 2025).

## 5. Conclusions

Wet sanitation in LMF production facilities can unintentionally promote pathogen growth. Elevated RH during and after sanitation slows the drying of soiled surfaces, keeping the water *a*_w_ of residues above the minimum threshold required for pathogen growth. Under these conditions, *Salmonella*, *Listeria monocytogenes*, and *Cronobacter sakazakii* can proliferate on surfaces, increasing the risk of cross-contamination.

Current sanitation guidance for LMF production environments emphasize minimizing direct water use and ensuring complete surface drying yet control of ambient RH is rarely addressed. This study demonstrated that LMF safety guidelines that do not address RH are incomplete, as they fail to account for the role of ambient RH in sustaining conditions conducive to pathogen proliferation. Incorporating RH control and monitoring in LMF safety programs would be a proactive means of preventing environmental microbial growth, thereby reducing the likelihood of cross-contamination events.

